# SCIITensor: A tensor decomposition based algorithm to construct actionable TME modules with spatially resolved intercellular communications

**DOI:** 10.1101/2024.05.21.595103

**Authors:** Huaqiang Huang, Chuandong Liu, Xin Liu, Jingyi Tian, Feng Xi, Mei Li, Guibo Li, Ao Chen, Xun Xu, Sha Liao, Jiajun Zhang, Xing Liu

## Abstract

Advanced spatial transcriptomics (ST) technology has paved the way for elucidating the spatial architecture of the tumor microenvironment (TME) from multiple perspectives. However, available tools only focus on the static molecular and cellular composition of the TME when analyzing the high-throughput ST data, neglecting to uncover the in-depth spatial co-variation of intercellular communications arising from heterogeneous spatial TMEs. Here, we introduce SCIITensor, which decomposes TME modules from the perspective of spatially resolved intercellular communication by spatially quantifying the cellular and molecular interaction intensities between proximal cells within each domain. It then constructs a three-dimensional matrix, formulating the task as a matrix decomposition problem, and identifies biologically relevant spatial interactions and TME patterns using Non-Negative Tucker Decomposition (NTD). We evaluated the performance of SCIITensor on liver cancer datasets obtained from multiple ST platforms. At the research setting of a single-sample investigation, SCIITensor precisely identified a functional TME module indicating a tumor boundary structure specific domain with co-variant interaction contexts, which were involved in construction of immunosuppressive TME. Moreover, we also proved that SCIITensor was able to construct TME meta-modules across multiple samples and to further identify an immune-infiltration associated and sample-common meta-module. We demonstrate that SCIITensor is applicable for dissecting TME modules from a new perspective by constructing spatial interaction contexts using ST datasets of individual and multiple samples, providing new insights into tumor research and potential therapeutic targets.

## Introduction

The tumor microenvironment (TME) is a complex and coordinated multicellular ecosystem with features including immune cells, stromal cells, blood vessels, and extracellular matrix (ECM)^1^. It is not just a simple composition of these components, but rather an actively connected system with bidirectional intercellular and intermolecular interactions that play a decisive role in carcinogenesis, tumor progression, metastasis, recurrence, and therapeutic response^1,2^. Systematically dissecting the spatial interaction context of the TME will enhance our comprehensive understanding of tumors and facilitate the next generation of treatment strategies^3^. Single-cell RNA sequencing (scRNA-seq) has ushered in a new epoch of research on the TME in recent years^4–6^. However, the tissue dissociation procedure of scRNA-seq results in the loss of spatial information of cells, hindering the exploration of complex organ-like structures with specific localized cellular phenotype distribution and spatially dynamic interactions in the TME^7^.

Recently, advances in spatially resolved transcriptomics have enabled the in-situ deconstruction of TME^8^. Several computational tools have been developed to analyze the TME using ST datasets^9–14^. Most of the tools analyze the cellular composition of the TME, such as CytoCommunity^9^, CC (local cell composure)^10^, CF-IDF^11^, and Spatial-LDA^12^. These tools resolve the cellular phenotype construction in each TME unit or the nearby cellular frequencies of each cell, and then cluster them into Cellular Neighborhood (CN) communities using graph-based community detection, k-means, or natural language processing methods. There are other tools designed to infer spatially proximal cell-cell communication by leveraging the spatial information of ST data, such as SpaTalk^13^, CellChat v2^14^, and others^15–17^. However, no method has been developed to systematically and quantitatively dissect the TME from the perspective of spatial interactions among cells and molecules.

Within this context, we present SCIITensor, a framework that decomposes the patterns of TME units and the spatial interaction modules based on NTD^18^, an unsupervised method that can identify spatial patterns and modules from multi-dimensional matrices with good interpretability. SCIITensor constructs a three-dimensional matrix by stacking intensity matrices of interactions in each TME unit, and it is decomposed by NTD. The decomposed patterns in each dimension indicate events related to specific cellular and molecular function modules within particular TME modules. Matrices of multiple samples can continue to be stacked together after replacing their third dimension with decomposed TME patterns. Factorizing the extended three-dimensional matrix can reveal meta-modules of TME units across multiple samples.

To illustrate the performance of our approach, we utilized SCIITensor to analyze Stereo-seq datasets of liver cancer. The result of individual sample uncovered an immunosuppressive TME module characterized by critical interactions between hepatocytes and macrophages, which have been confirmed in a prior study^19^, at the tumor border of intrahepatic cholangiocarcinoma (ICC). The result of multiple samples uncovered an infiltration TME meta-module characterized by interactions between fibroblasts/Malignant cells and macrophages. We also successfully applied SCIITensor to low-throughput ST data obtained by MERFISH. Collectively, SCIITensor is a broadly applicable method that can systematically dissect TME modules from a new perspective and uncover spatially resolved interaction patterns for both individual and multiple samples, providing new insights into the biologically meaningful structure and function of tumors.

## Results

### Overview of SCIITensor

Overall speaking, SCIITensor deciphered the TME patterns from the perspective of spatial interactions in an unsupervised manner (Fig. 1). It consists of three components: (1) constructing a three-dimensional interaction matrix based on calculating spatial cell interaction intensity (SCII)^20^ in each TME unit, (2) decomposing the TME and spatial interaction modules in individual samples by factorizing the constructed interaction tensor, and (3) constructing the meta-modules of TME across multiple samples by factorizing the merged interaction tensor. Meanwhile, provide spatial interaction modules which are associated with these TME modules and meta-modules.

**Fig 1.**
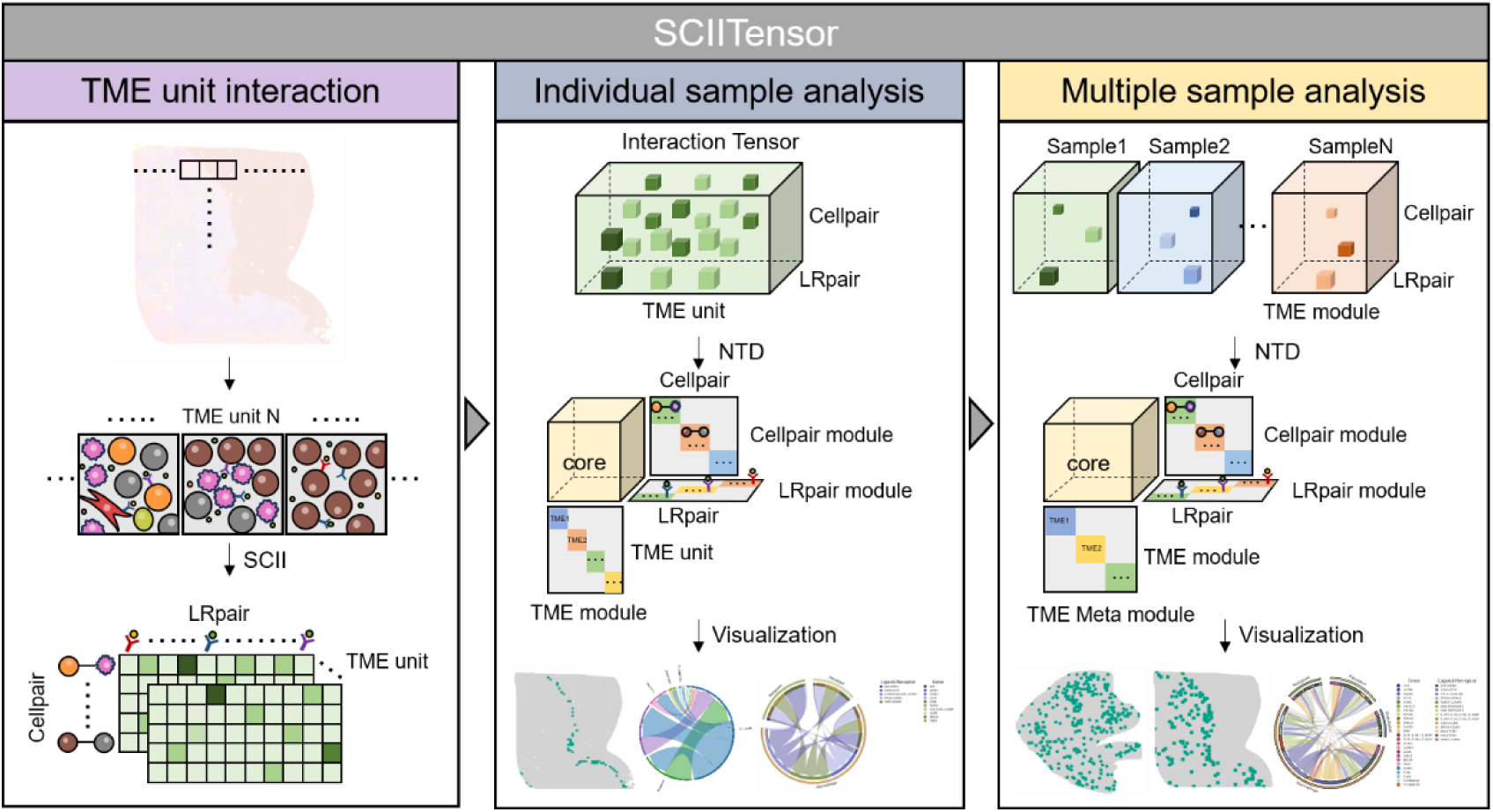
Overview of the SCIITensor algorithm. SCIITensor first divides ST data into TME units using adjacent square windows. Then, it estimates the spatial cellular interaction intensity in each TME unit to generate interaction matrices with dimensions of cell pairs and LR pairs. Next, these interaction matrices are integrated into a three-order interaction tensor. Non-negative Tucker decomposition is then applied to decompose interaction modules and TME modules in individual samples. The identified TME modules are shown in situ, and the cellular and molecular interactions strongly correlated to it are displayed using circle plots. The identification of TME meta-modules across multiple samples is supported. Firstly, stack the transformed matrices in which the third dimension TME units have been replaced by decomposed TME modules of each sample. Then, NTD is employed to factorize the merged interaction tensor to investigate TME meta-modules, CC modules and LR modules. The TME meta-modules are plotted in situ, and interactions associated with them are illustrated using circle plots.

SCIITensor divides the ST-sequenced tissue region into TME units using adjacent square windows. Each TME unit is characterized by profiling of ligand-receptor molecules mediating communication between different phenotypic cells. To investigate the patterns of the TME organized by similar spatial interactions, we construct the spatial neighborhood interaction graph using SCII. This method enables the quantitative measurement of spatially resolved interactions between neighboring cells within each TME unit. Subsequently, the interaction matrices obtained, with dimensions of cell pairs and ligand-receptor (LR) pairs for each TME unit, were combined into a three-order interaction tensor. The non-negative scores of the interaction tensor represent the strength of interactions between cells within the TME units.

Next, NTD^18^ was implemented to factorize the constructed three-dimensional interaction tensor into a core matrix and three low-dimensional factor matrices. The decomposed factor matrices represent modules of cell pairs, LR pairs, and TME units, while the core matrix represents the relationships across all the different modules. The decomposed results can reveal TME modules structured by specific cell-cell communications mediated by particular LR molecules, identifying biologically meaningful patterns, structures, and tissue regions in situ. The relationship between TME modules and cellular and molecular interaction modules can be quantified by the scores in the core matrix.

In order to analyze the meta-patterns of TME across multiple samples, SCIITensor stacked the transformed spatial interaction tensors with dimensions of cell pairs ×LR pairs ×TME modules. The TME modules were obtained from the factorization result of the original tensor. Then we decoded the TME and spatial interaction patterns across multiple samples by applying NTD to the merged tensor. We can obtain sample-specific and sample-common TME meta-modules while identifying the cell-cell communication features within these meta-modules. To visualize the results, SCIITensor implemented several visualization methods, including circle plots, heatmaps, and Sankey diagrams to assist researchers select L-R of interest within selected module.

### SCIITensor distinguishes cell communication defined TME modules of liver cancer

To demonstrate the capabilities of the SCIITensor, it was used to reveal an immunosuppressive TME module located around the tumor border from a ST dataset of ICC^19^ (Fig. 2). Firstly, we utilized cell2location^21^ to deconvolute the Stereo-seq data in bin14 (∼10um) spatial enhanced resolution. Each binned spot was treated as a cell and labeled according to the cell type with the highest proportion (Fig. 2A). The accuracy of the cell annotation result was validated by the higher expression of classical cell type marker genes (Fig. S1A). Next, SCIITensor was applied to construct TME modules from the perspective of cellular and molecular interactions. To determine the optimal rank combination for the factorization of the interaction tensor, we explored a range of rank combinations and identified that the combination of 15, 15, and 14 resulted in the lowest reconstruction error (Fig. 2B).

**Fig 2.**
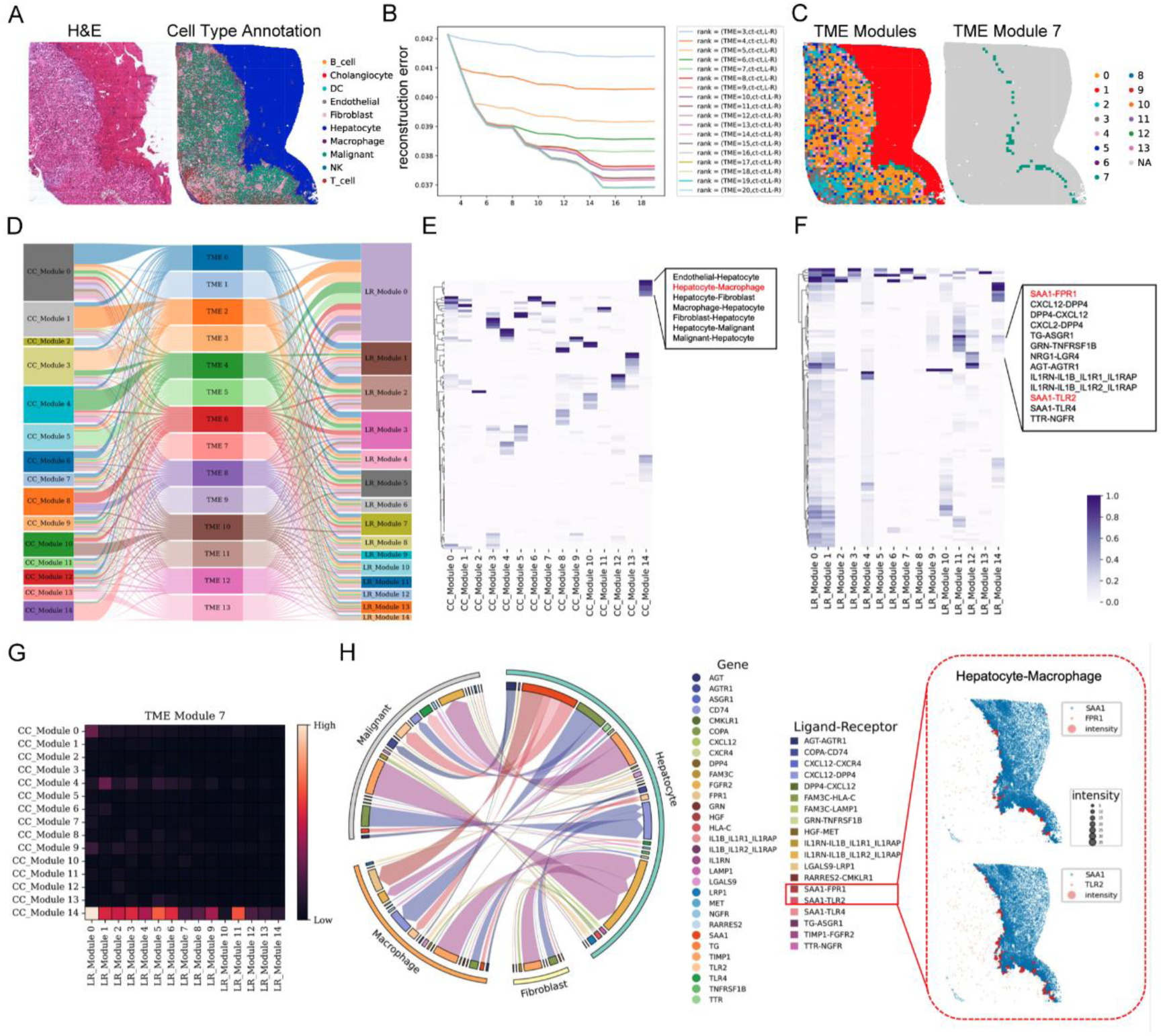
Unraveling the TME module located at the tumor border of liver cancer. **A**. H&E staining (left) and cell annotation (right) of LC5M sample. **B.** The line graph shows the reconstruction errors for LC5M with various rank combinations. The colors of the lines indicate the ranks for the decomposition of TME units. The x-axes represent the ranks of CC-pairs and LR-pairs, which have been set to be the same, while the y-axes represent reconstruction errors. **C.** In situ display of distinct TME modules (left) and TME module 7 (right). **D.** Sankey diagram illustrates the relationship between TME modules, CC modules, and LR pair modules. **E, F.** Heatmaps represent the factor matrices of the CC module and LR module. The colors indicate the contribution of CC pairs or LR pairs to corresponding CC modules or LR modules. **G.** The heatmap indicates the contributions of CC modules and LR modules to TME module 7. The colors represent the strength. **H.** The circular plot (left) shows interactions that significantly contribute to TME module 7. The spatial plots (right) show interactions in situ between hepatocytes and macrophages mediated by SAA1-FPR1 and SAA1-TLR2. Blue dots represent hepatocytes expressing SAA1, while orange dots represent macrophages expressing FPR1 or TLR2. Red dots represent interactions between hepatocytes and macrophages mediated by SAA1-FPR1/TLR2, with the size indicating the intensity of interactions. CC: Cell-Cell, LR: Ligand-Receptor.

The TME units were clustered into 14 TME modules, and TME module 7 was located around the tumor margins (Fig. 2C). Tumor margins are regions with highly active communications among malignant cells, hepatocytes and macrophages, along with comprehensive tumor cell infiltration and invasion^19^. To explore the active cellular and molecular architecture in the crucial region where TME module 7 was located, we visualized the decomposed core matrix, which represented the relationships among TME modules, cell pair modules, and LR pair modules (Fig. 2D). The associations of cell pair modules and LR pair modules with TME module 7 were displayed using a heatmap (Fig. 2G). Strong relationships were observed between TME module 7 and CC module 14, as well as LR modules 0, 5, and 11. Furthermore, the factor matrices representing the contributions of specific cell pairs or LR pairs to the corresponding CC modules or LR modules were visualized (Fig. 2E, F). The hepatocyte-macrophage pair was identified to highly contribute to CC module 14, while the SAA1-FPR1/TLR2 pairs highly contributed to LR module 11. This suggests the possible molecular activity in which SAA1 secreted by hepatocytes interacts with FPR1 and TLR2 of macrophages in the TME module 7 (Fig. 2H, Fig. S1B). This finding is consistent with the original study^19^. The SAA1-FPR1 and SAA1-TLR2 have been reported to participate in the recruitment of macrophages and M2 polarization, thereby promoting the establishment of a local immunosuppressive microenvironment^19^. Collectively, SCIITensor successfully identified interpretable TME modules from the perspective of communication architectures. The verified immunosuppressive TME region around the tumor border has been unsupervised identified without errors introduced by manual selection. At the same time, the communication context in this TME module has been correctly and clearly decoded.

### Identification of infiltration TME meta-module across multiple samples using SCIITensor

Investigation of sample-common and sample-unique TME modules across multiple samples is critical for tumor research. Integrating samples from multiple regional sites to detect the sample-common TME modules can help to understand the complex biological behavior of tumors from a broader perspective. Similarly, integrating samples from different conditions, such as treatment versus control, to detect the sample-specific TME modules can aid in investigating the response or resistance mechanisms of tumor treatment. To address the requirement, we developed the multiple sample analysis module of SCIITensor and applied it to analyze three samples obtained from different anatomical sites of the same liver cancer patient, including tumor core (LC5T), tumor margin (LC5M), and paratumor tissue (LC5P)^19^. The cell type annotation was performed using cell2location, and the higher expression levels of classical marker genes of corresponding cell types verified the accuracy of the annotation (Fig. S2A, B). The combination of ranks 14, 14, and 17 with the lowest reconstruction error was selected to decompose sample LC5T (Fig. 3A), while the combination of ranks 16, 16, and 13 with the lowest reconstruction error was selected to decompose sample LC5P (Fig. 3B). The decomposed result of sample LC5M has been obtained in the above figure (Fig. 2B, C). After replacing the third dimension of the interaction tensors with corresponding decomposed TME modules, they are stacked together into an integrated interaction tensor. Subsequently, the combination of ranks 15, 15, and 25 with the lowest reconstruction error was selected for decomposition (Fig. S2C).

**Fig 3.**
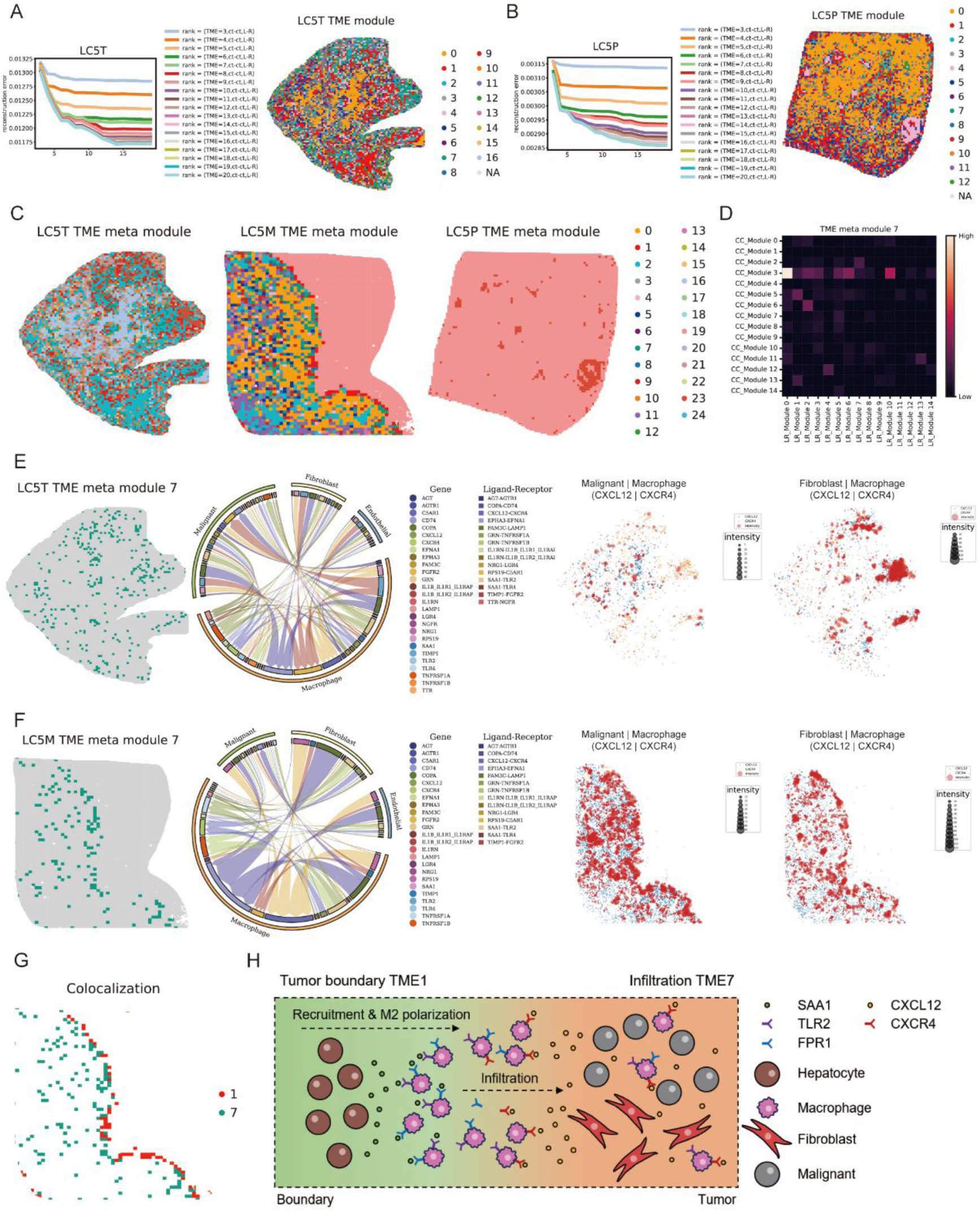
Reveal TME meta-modules across multiple samples obtained from different regional sites of a liver cancer patient. **A**. The line graph (left) shows the reconstruction errors for LC5T with different combinations of ranks. The colors represent different ranks of TME units. Spatial distribution (right) of TME modules of LC5T. **B.** The line graph (left) shows the reconstruction errors for LC5P with different combinations of ranks. The colors represent different ranks of TME units. Spatial distribution (right) of TME modules of LC5P. **C.** Spatial distribution of TME meta-modules of LC5T, LC5M, and LC5P, respectively. **D.** The heatmap indicates the contribution of CC modules and LR modules to TME meta module 7. The colors represent strength. **E, F.** Spatial distribution (left) of TME meta-module 7 in LC5T or LC5M; Circle plot (middle) displays interactions that significantly contribute to TME meta-module 7. Spatial interactions (right) between malignant/fibroblast cells and macrophages mediated by CXCL12-CXCR4 are shown in situ. Blue dots represent malignant cells with CXCL12 expression, while orange dots represent macrophages with CXCR4 expression. Red dots indicate interactions between malignant/fibroblast cells and macrophages mediated by CXCL12-CXCR4, with the size of the red dots reflecting the intensity of the interaction. **G.** Spatial distribution of TME meta-modules 1 and 7. **H.** Schematic diagram shows that hepatocytes secrete SAA1 to recruit FRP1+/TLR2+ macrophages and polarize them into the M2-like phenotype at the tumor border, then the macrophages expressing CXCR4 infiltrate to the tumor core by the attraction of CXCL12 secreted by malignant cells/fibroblasts.

The TME units of multiple samples were clustered into 25 TME meta-modules across them. There are some sample-specific TME meta-modules, such as TME meta-modules 10, 16, 18, 20, 21, 22, 24 uniquely distributed in LC5T, TME meta-modules 0, 1, 3, 4, 5, 8, 11, 12, 13 uniquely distributed in LC5M, while other TME meta-modules are sample-common, such as TME meta-modules 2, 7, 9 distributed in both LC5T and LC5M, TME meta-module 23 distributed in both LC5T and LC5P, and TME meta-module 19 distributed in both LC5M and LC5P (Fig. 3C, S2G). To be noted, we found that the edge of TME meta-module 7 in LC5M closely colocalized with TME meta-module 1 (Fig. S2G), which is consistent with the TME module 7 in individual sample analysis of LC5M. Meta-module 1 is a unique module structuring the tumor boundary, which is observed solely in para-tumor sample (LC5M). Meta-module 7 is a sample-common module observed both in tumor core sample and para-tumor sample. To determine the underlying functional correlation between TME meta-module 7 and 1, we extracted the interaction information of TME meta-module 7 and visualized the relationship between TME meta-module 7 and CC modules, as well as LR modules, using a heatmap (Fig. 3D). Subsequently, the cellular and molecular communications of CC module 3 and LR module 11, which contributed more strongly to TME meta-module 7, were visualized using a circle diagram (Fig. 3E, F). The interactions between malignant cells/fibroblasts and macrophages mediated by RPS19-C5AR1, COPA-CD74, and CXCL12-CXCR4 greatly contributed to the architecture of TME meta-module 7.

CXCL12-CXCR4 has been reported to play a critical role in monocyte infiltration and polarization of macrophages to the M2-like phenotype^22,23^, which may contribute to the formation of a local immunosuppressive microenvironment. In order to investigate the specific spatial interactions facilitated by CXCL12-CXCR4 between malignant cells/fibroblasts and macrophages in various tissue regions, they were visualized in situ (Fig. 3E, F, S2H). The overall expression levels of these LR molecules in different cell types were depicted using a bubble plot (Fig. S2I). Compared with the tumor border (LC5M), communications between malignant cells/fibroblasts and macrophages mediated by CXCL12-CXXR4 were weaker in the tumor center (LC5T), suggesting a decreasing gradient of macrophage infiltration from the tumor border to the tumor center. Integrating the interesting finding of TME meta-module 7 with the previous finding of TME meta-module 1, we imply that TLR2+/FRP1+ macrophages in TME meta-module 1 were recruited by the SAA1+ hepatocytes and polarized to an M2-like phenotype. Subsequently, the M2-like macrophages express CXCR4 and infiltrate the tumor due to the attraction of CXCL12 secreted by malignant or fibroblast cells, as observed in TME meta-module 7 (Fig. 3G, H). Taken together, the integrated analysis results validate the capability of SCIITensor to identify TME meta-modules and decode the comprehensive biological meaningful mechanism across multiple samples.

### The performance of SCIITensor in MERFISH data

In addition to sequencing-based high-throughput ST technologies like Stereo-seq^24^, there are also image-based low-throughput ST technologies such as MERFISH^25^. To investigate the capability of SCIITensor to decipher low-throughput ST data, we applied it to MERFISH data from liver cancer. To annotate the ST data, we clustered cells and assigned a cell type label to each cluster based on their highly expressed genes (Fig. 4A). The interaction tensor was then constructed and decomposed using the optimal combination of ranks 4, 4, and 3 (Fig. S3B). Three TME modules with distinct cellular and molecular interaction contexts, four CC modules, and four LR modules were distinguished in the liver cancer sample (Fig. 4B-E).

**Fig 4.**
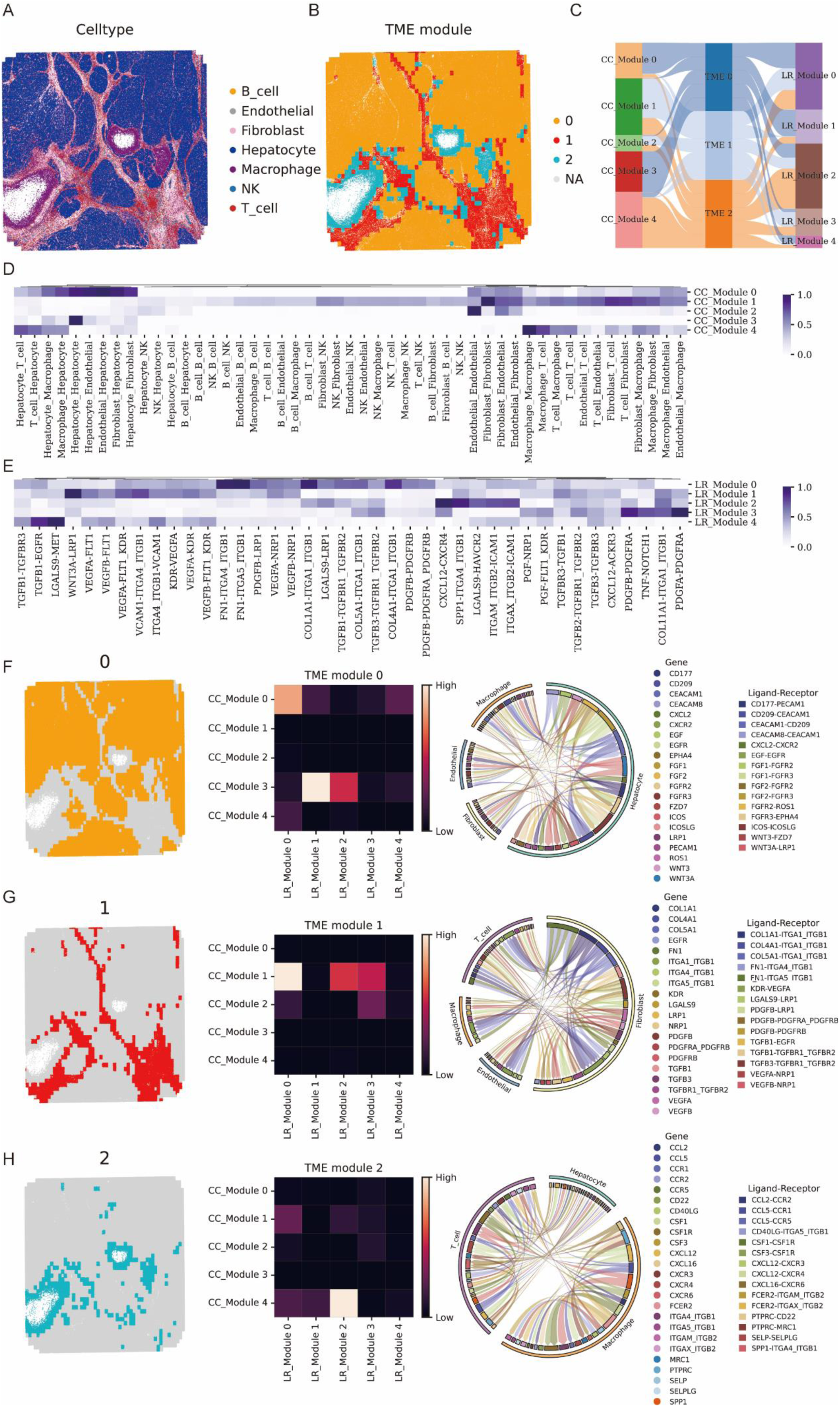
SCIITensor detects TME patterns and their corresponding communication architectures in a liver cancer dataset obtained by MERHISH. **A**. Spatial distribution of annotated cells. **B.** The spatial distribution of TME modules. **C.** Sankey diagram illustrates the correlation among CC modules, LR modules, and TME modules. **D, E.** Heatmaps represent the CC module factor matrix and the LR module factor matrix, where colors indicate the strength of contribution of CC pairs or LR pairs to each module. **F, G, H.** The spatial distribution (left) of TME modules 0, 1, and 2; Heatmaps (middle) indicate the strength of relationships between CC modules, LR modules, and their corresponding TME modules; The circle plot (right) shows interactions that strongly contribute to TME modules 0, 1, and 2.

The TME module 0, which was more strongly correlated with CC module 3 and LR module 1, was characterized by interactions among stromal cells (fibroblasts, and endothelial cells), hepatocytes and macrophages mediated by fibroblast growth factors (FGF1, FGF2) and FGF receptors, as well as Wnt ligands and their receptors (FZD7, LRP1) (Fig. 4F and Fig. S3C). FGF1 and FGF2 are well-known for their angiogenic properties as they induce the promotion of endothelial cell proliferation and the physical organization of endothelial cells into tube-like structures^26^. Wnt ligands have been investigated to activate the Wnt/β-catenin pathway through FZD7 in HCC cells and may play a role during hepatocarcinogenesis^27^. Both WNT3 and WNT3A are independent prognostic factors for hepatocellular carcinoma (HCC)^28,29^, and silencing WNT3A inhibits the growth of liver cancer^30^. In TME module 0, it is suggested that activated hepatocytes are malignant, and progression is promoted through specific active interactions. The TME module 1, which exhibited a stronger correlation with CC module 1 and LR module 0, was characterized by interactions between fibroblasts mediated by cell adhesion-associated LR pairs such as FN1-ITGA5_ITGB1, FN1-ITGA4_ITGB1, and COLIA1-ITGA1_ITGB1, and angiogenesis-associated^31^ LR pair VEGFA-NRP1 (Fig. 4G and Fig. S3D). The LR pairs involving fibronectin (FN1) and collagen (COL1A1) suggest that fibroblasts may be involved in remodeling the extracellular matrix (ECM) in TME module 1 as fibronectin and collagen are major components of the ECM and play crucial roles in cell adhesion and migration. The interaction between fibroblasts and these ECM proteins may contribute to tumor cell invasion and metastasis. The presence of VEGFA-NRP1, which is associated with angiogenesis, suggests that fibroblasts may promote angiogenesis in the TME module 1. In TME module 2, which were more strongly correlated with CC module 4 and LR module 2, interactions between macrophages mediated by CXCL12-CXCR4/CXCR5, CCL5-CCR1/CCR5, and CSF1/CSF3-CSF1R were identified (Fig. 4H and Fig. S3E). The interactions between chemokines (CXCL12, CCL5) and their corresponding receptors (CXCR4/CXCR5, CCR1/CCR5) indicate that macrophages are recruited and attracted to the locations of the TME module 2 and are involved in the inflammatory response as chemokines like CCL5 are known to play a role in immune cell recruitment and activation^22^. The interactions involving colony-stimulating factors (CSF1/CSF3) and their receptor (CSF1R) have been verified to regulate macrophage differentiation and polarization^32^. Overall, SCIITensor successfully analyzed the low-throughput ST data.

## Discussion

Unlike previous approaches that investigate TME from the perspective of static molecular and cellular composition, SCIITensor is the first method for detecting TME modules that leverages the power of NTD to systematically distinguish TME modules from the perspective of active spatial cell interactions without supervision. Instead of detecting TME modules in individual samples, SCIITensor can be used to analyze TME meta-modules across multiple samples to investigate the complex sample-common or sample-specific TME meta-modules across different tissue regions. At the same time, SCIITensor can decipher CC modules, LR modules and crucial cellular and molecular interactions that shape various TME modules. Furthermore, the enhanced compatibility of SCIITensor with analyzing datasets generated by imaging-based low-throughput ST technologies such as MERFISH underscores its versatility and potential for widespread adoption. In principle, in addition to transcriptomics spatial data, it can analyze multiple omics spatial data such as spatial proteomics (Deep Visual Proteomics (DVP)^33^, DBiT-seq^34^).

Application of our SCIITensor method to datasets of liver cancer samples obtained from various ST technologies shows that it can successfully identify specific TME modules in individual samples, TME meta-modules across multiple samples, and the cellular and molecular communication context among these modules. The findings obtained by SCIITensor may have significant implications for understanding tumor biology and identifying potential therapeutic targets in cancer treatment.

Due to the necessity of constructing spatial cell interaction graphs, the current version of SCIITensor is unsuitable for spatial omics datasets with low resolution. To address this issue, spatial cell communication can be inferred from deconvoluted cellular composition while constructing spatial molecular interaction graphs within each TME unit. Then, integrate the cellular and molecular information to detect TME modules.

In conclusion, SCIITensor opens new avenues for a systematic understanding of the TME by examining spatial cellular and molecular interactions. As the rapid growth of spatial omics data at single-cell resolution continues, we anticipate that SCIITensor will be a powerful and scalable method for investigating TME, providing deep insights in cancer research.

## Methods

### Data preprocessing and cell type annotation

*Single-cell RNA data*. Preprocessing was performed using the Seurat^35^ package. Briefly, cells with nFeature_RNA ranging from 500 to 6000, percent.mt below 20, and nCount_RNA exceeding 2000 were included in subsequent analyses. Then normalization, selection of highly variable genes, standardization, principal component analysis (PCA), and uniform manifold approximation and projection (UMAP) were used to reduce dimensionality and visualize the data. Leiden clustering was applied to differentiate cell types with curated markers. FXYD2, TM4SF4, and ANXA4 are specific markers for cholangiocytes. APOC3, FABP1, and APOA1 are specific markers for hepatocytes. MS4A1 and CD79A are specific markers for B cells. CD2, CD3D, and CD3E are specific markers for T cells. CD14, CHIT1, CYP19A1, HK3, MSR1, VSIG4, CLEC5A, ADAMDEC1, and DNASE2B are specific markers for macrophages. CD7, FGFBP2, KLRF1, and NCAM1 are specific markers for natural killer (NK) cells. CLEC9A and CD1C are specific markers for dendritic cells (DC). ACTA2 and COL1A2 are specific markers for fibroblasts. CLDN5, PECAM1, CD34, FLT1, VWF, ENG, and CDH5 are specific markers for endothelial cells.

*Stereo-seq data*. The stereo-seq data of the ICC was divided into adjacent windows, each with an area of 20×20 DNBs (bin20). Genes detected in fewer than three bin20 units were filtered out, while bin20 units with fewer than three detected genes were excluded from the data analysis. The cell type composition of each bin20 was inferred using the cell2location^21^ package with ICC scRNA-seq data as a reference. The highest-ranked cell type was assigned to each spot as its cellular identity.

*MERFISH data*. Preprocessing was conducted using the Scanpy^36^ package. Cells with fewer than 10 detected RNA counts were excluded. Then, normalization, selection of highly variable genes, standardization, PCA, UMAP, and Leiden clustering were employed. The cell type of each cluster was determined based on the expression of the top four differentially expressed genes.

## SCIITensor

### Construction of the spatial cell-cell interaction tensor

We combined 10×10 neighboring bin20 units into 200×200 windows (TME units), meaning each window contains 10×10=100 bin20 units. Then, spatial neighborhood graphs were generated for each LR pair within each TME unit, using a neighborhood radius threshold of 200 DNB. This means that only the sender bin20 was connected to the receiver bin20 units whose distance is less than 200 DNB, and the corresponding ligand and receptor genes are expressed. The weight of each edge linked between bin20 units is equal to the sum of the ligand expression in the sender bin20 and the receptor expression in the receiver bin20. The Spatial Cellular Interaction Intensity (SCII)^20^ between each cell type was calculated, depicting the patterns of cell-cell communication in each TME unit. The SCII is calculated as follows:

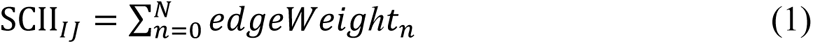

Where: *I* is the sender cell type, *J* is the receiver cell type, *n* is n-th edge that link sender cell *i* and receiver cell *j*. *N* represents the number of edges that link cell type *I* and cell type *J*.

Finally, we combined all TME unit interaction matrices into a third-order interaction tensor *X* ∈ *ℝ*^*O*×*L*×*S*^ with dimensions cell pairs ×LR pairs ×TME units. Cell pairs, LR pairs, and TME units with all zero values in the interaction tensor were excluded from subsequent analysis, then the interaction tensor was log-transformed.

### Identification of communication and TME patterns in individual samples

To decipher the interaction and TME patterns of individual samples, SCIITensor employs non-negative tucker decomposition (NTD)^18^ implemented in the Tensorly^37^ Python package on the three-dimensional interaction tensor *X* ∈ *ℝ*^*O*×*L*×*S*^, factorizing it into a core matrix and three matrices as:

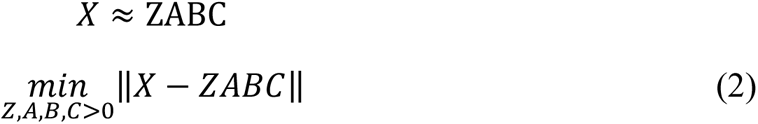

Where the three low-dimensional matrices *A* ∈ *ℝ*^*O*×*P*^, *B* ∈ *ℝ*^*L*×*Q*^, *C* ∈ *ℝ*^*S*×*R*^ are the cell pair loading, LR pair loading and TME unit loading factor matrices with size *O* × *P*, *L* × *Q* and *S* × *R*, respectively. These factor matrices capture the unique patterns and variations along each dimension. *A* ∈ *ℝ*^*O*×*P*^represents the relationships and patterns along the first dimension (cell pairs) of the original matrix *X*, *A*_*o*×*p*_ is the loading values of cell pair *o* in pattern *p*, representing the contributions of cell pair *o* in pattern *p*. *B* ∈ *ℝ*^*L*×*Q*^ represents the relationships and patterns along the second dimension (LR pairs) of the original matrix *X*, *B*_*l*×*q*_is the loading values of LR pair *l* in pattern *q*, representing the contributions of LR pair *l* in pattern *q*. *C* ∈ *ℝ*^*S*×*R*^represents the relationships and patterns along the third dimension (TME units or 200×200 windows), *C*_*s*×*r*_ is the loading values of TME unit *s* in pattern *r*, representing the contribution of TME unit *s* in pattern *r*. *Z* ∈ *ℝ*^*P*×*Q*×*R*^is the core matrix compactly representing the original three-dimensional matrix with size *P* × *Q* × *R*. It contains shared patterns and relationships across all dimensions. The entries in the core matrix represent the strengths or contributions of these patterns.

In summary, factor matrices can offer insights into the underlying patterns and relationships within each dimension, while the core matrix represents the shared patterns across all dimensions. The matrix *A* represents the *P* latent patterns of cell pair groups, indicating how these cell pair groups coordinate to interact. The matrix *B* represents the *Q* latent patterns of ligand-receptor pairs, indicating how these pairs work to communicate. The matrix *C* represents the *R* latent patterns of TME units, indicating which TME units have similar communication patterns. The core matrix *Z* represents the connections between patterns in three dimensions, predicting the key ligand-receptor pairs drive the communication between specific sender-receiver cell pairs, and the key sender-receiver cell pairs and corresponding ligand-receptor pairs construct specific TME.

To determine the optimal rank for NTD, SCIITensor decomposes the three-dimensional matrix *X* using a range of possible rank combinations and calculates their reconstruction error by comparing the original three-dimensional matrix with the reconstructed matrix. Then plot the reconstruction error for each rank combination on a graph. To reduce the computational complexity, the rank of cell pairs and LR pairs dimensions are set to be equal. Look for the visual elbow point at which the error starts to plateau or decrease at a slower rate. The rank combination corresponding to the point of diminishing returns in the reconstruction error will give us a good balance between accuracy and complexity.

To intuitively show the associations of latent patterns with cell pairs, LR pairs, and TME units, we used Sankey diagrams and heatmaps. The core matrix was visualized as a Sankey diagram using the matplotlib^38^ Python package. The connection information between the TME module of interest and its corresponding cell pair modules and ligand-receptor modules was extracted from the core matrix. The extracted connection matrix of the TME module of interest was then displayed as a heatmap diagram using the seaborn^39^ Python package. Based on the matrix data, the top-ranked and interested cell pair module and LR module that strongly contribute to the TME module of interest were selected. Then, the top 5 cell pairs in the selected cell pair module and the top 15 LR pairs in the selected LR module were visualized using a circle plot generated by the pycirclize Python package and a heatmap created with the seaborn Python package. Specific cell-cell interactions were visualized in situ using the StereoSiTE Python package.

### Identification of communication and TME meta-patterns in multiple samples

To investigate the pattern of cell-cell communication across multiple samples, we developed the multiple sample analysis workflow of SCIITensor. Firstly, the interaction tensor *X* ∈ *ℝ*^*O*×*L*×*S*^ of each individual sample was decomposed using the communication and TME pattern identification function of SCIITensor mentioned above. Secondly, based on the parsed factor matrix *C* ∈ *ℝ*^*S*×*R*^, indicating connections between TME units with TME modules, the third dimension *S* representing TME units of the tensor *X* ∈ *ℝ*^*O*×*L*×*S*^was transferred to the TME modules with size *R* by the operation below:

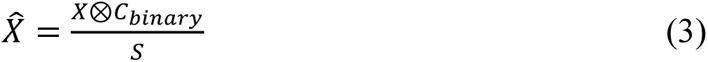

Where: 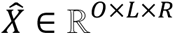 is the transformed interaction tensor with dimensions *O* × *L* × R; *X* ∈ *ℝ*^*O*×*L*×*S*^ is the original interaction tensor with dimensions *O* × *L* × *S*; *C*_*binary*_ ∈

*ℝ*^*S*×*R*^ is the factor matrix of TME module with dimensions *S* × *R*. For each *S* dimension, the maximal value was set to 1 and others was set to 0. *S* denotes the number of the TME units.

To perform this matrix operation, we performed element-wise multiplication between the 2D factor matrix *C*_*binary*_and the corresponding elements of the 3D interaction matrix *X*. This multiplication was done for each combination of *O*, *L*, and *R* indices:

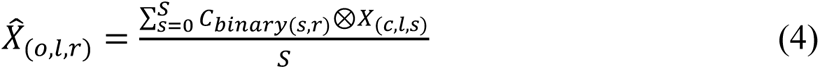

Where: 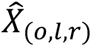 represents the element of 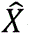 at the o-th row, l-th column, and r-th depth. *C*_*binary*(*s*,*r*)_ represents the element of *C* at the s-th row and r-th column. *X*_(*c*,*l*,*s*)_ represents the element of *X* at the c-th row, l-th column, and s-th depth.

To identify the interaction structure across multiple samples, we integrated all transformed tensors 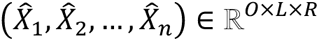 into a third-order tensor 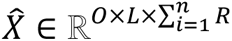 with dimensions cell pairs ×LR pairs ×TME modules. Then, NTD was performed to decompose the third-order tensor:

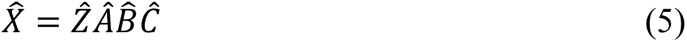

Where the three low-dimensional matrices 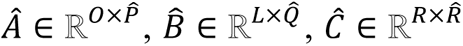 are the cell pair loading, LR pair loading and TME module loading factor matrices with size 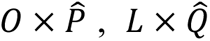 and 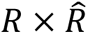, respectively. 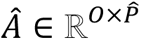 and 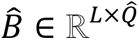 represent the relationship between cell pairs and LR pairs with their patterns as described in the individual sample NTD, 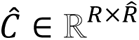 represents the relationship between TME modules of individual sample with meta TME modules across all samples. 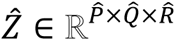 denotes the core matrix, representing the strength of the relationship between cell pair modules, LR pair modules and meta TME modules.

The determination of the optimal rank combination and visualization of decomposition results were performed according to the same schedule described in the individual sample analysis.

## Data availability

The scRNA-seq data of the ICC was obtained from the Gene Expression Omnibus (GEO) under accession number GSE138709. The stereo-seq data of the ICC was downloaded from the China National GeneBank (CNGB) Sequence Archive under accession number CNP0002199. The MERFISH data of liver cancer is available in the MERSCOPE FFPE Human Immuno-oncology Data Release (https://info.vizgen.com/ffpe-showcase, liver cancer 1).

## Code availability

The Python package SCIITensor, developed in this study, is publicly available on Github (https://github.com/STOmics/SCIITensor).

## Acknowledge

We acknowledge members of China National GeneBank for assistance with data. This study is supported by Science and Technology Innovation Key R&D Program of Chongqing (CSTB2023TIAD-STX0002).

## Competing interests

The authors declare no competing interests.

**Fig S1.**
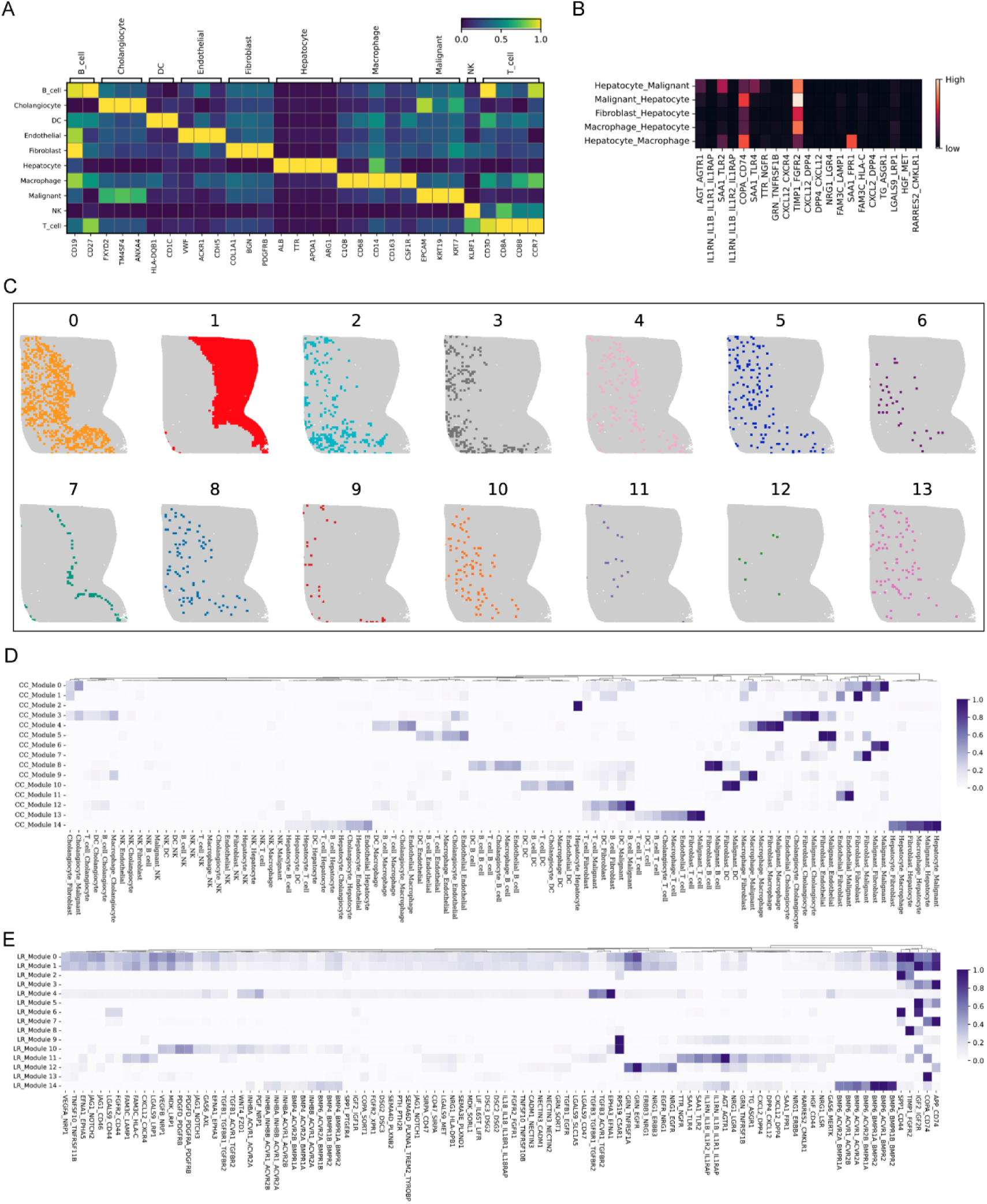
The analysis results of SCIITensor in LC5M. **A.** The heatmap indicates the expression levels of curated cell type markers. **B.** The heatmap shows the contribution strength of the top LR pairs and cell pairs to TME module 7. **C.** The spatial distribution of each TME module. **D.** The heatmap representing the CC module factor matrix. **E.** The heatmap displays the loadings of the top 120 LR pairs in the factor matrix.

**Fig S2.**
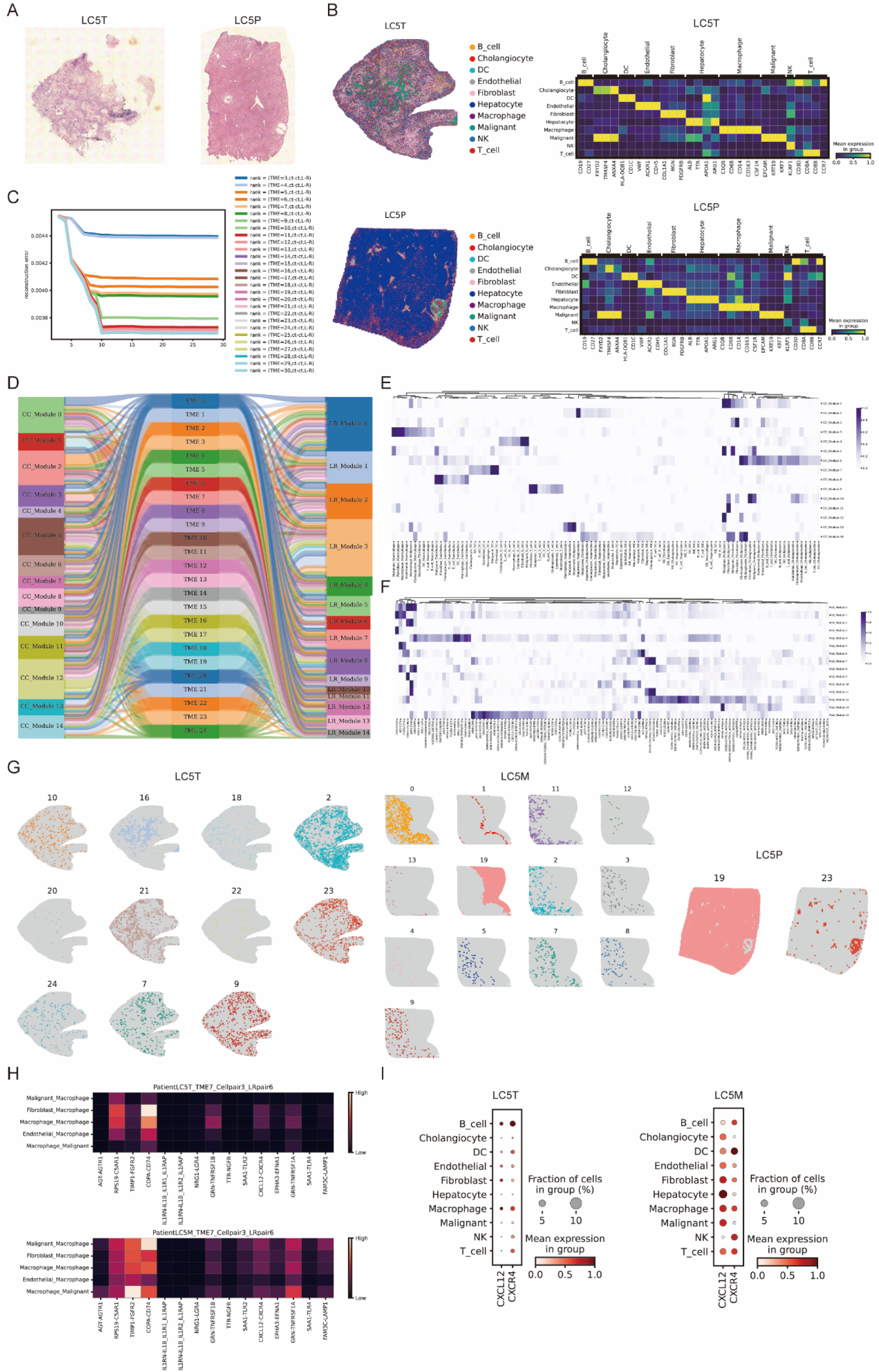
SCIITensor results across multiple samples include LC5T, LC5M, and LC5P. **A.** H&E staining of LC5T and LC5P. **B.** Spatial distribution of TME modules in LC5T (top left) and LC5P (bottom left). Heatmap (right) indicates the expression levels of curated cell type markers. **C.** The line graph displays the reconstruction errors corresponding to different rank combinations for decomposing the stacked interaction tensor across multiple samples (LC5T, LC5M, and LC5P). The colors represent different ranks of TME meta-modules. **D.** Sankey diagram illustrates the relationship between CC modules, LR modules, and TME modules. **E.** The heatmap indicates the factor matrix of the CC module. **F.** The heatmap shows the loadings of the top 120 LR pairs in the factor matrix. **G.** Spatial distribution of TME meta-modules in LC5T (left), LC5M (middle), and LC5P (right). **H.** The heatmap displays the contribution strength of the top LR pairs and cell pairs to TME meta module 7 in LC5T (top) and LC5M (bottom). **I.** Dot plots represent the expression levels of CXCL12 and CXCR4 in each cell type.

**Fig S3.**
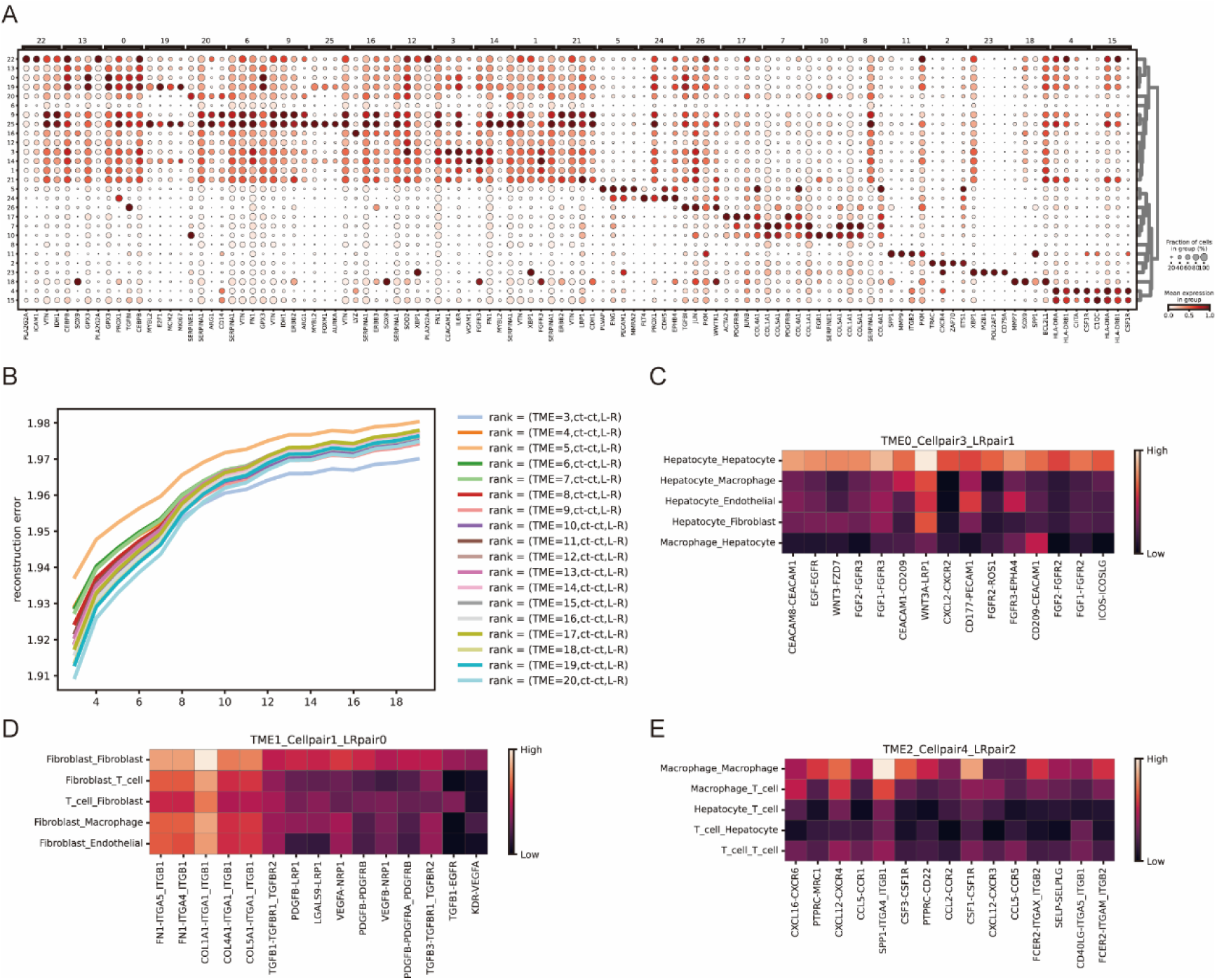
The results of SCIITensor in MERFISH data. **A.** Dot plot shows the expression levels of the top 4 differentially expressed genes in each cluster. **B.** The line graph displays the reconstruction errors corresponding to various combinations of ranks for decomposing the interaction tensor constructed from MERFISH data. The colors represent different ranks of TME modules. **C.** The heatmap displays the contribution strength of the top 15 LR pairs and the top 5 cell pairs to TME module 0. **D.** The heatmap displays the contribution strength of the top 15 LR pairs and the top 5 cell pairs to TME module 1. **E.** The heatmap displays the contribution strength of the top 15 LR pairs and the top 5 cell pairs to TME module 2.

## Notes

### Competing Interest Statement

The authors have declared no competing interest.

